# DrosoPhyla: genomic resources for drosophilid phylogeny and systematics

**DOI:** 10.1101/2021.03.23.436709

**Authors:** Cédric Finet, Victoria A. Kassner, Antonio B. Carvalho, Henry Chung, Jonathan P. Day, Stephanie Day, Emily K. Delaney, Francine C. De Ré, Héloïse D. Dufour, Eduardo Dupim, Hiroyuki F. Izumitani, Thaísa B. Gautério, Jessa Justen, Toru Katoh, Artyom Kopp, Shigeyuki Koshikawa, Ben Longdon, Elgion L. Loreto, Maria D. S. Nunes, Komal K. B. Raja, Mark Rebeiz, Michael G. Ritchie, Gayane Saakyan, Tanya Sneddon, Machiko Teramoto, Venera Tyukmaeva, Thyago Vanderlinde, Emily E. Wey, Thomas Werner, Thomas M. Williams, Lizandra J. Robe, Masanori J. Toda, Ferdinand Marlétaz

## Abstract

The vinegar fly *Drosophila melanogaster* is a pivotal model for invertebrate development, genetics, physiology, neuroscience, and disease. The whole family Drosophilidae, which contains over 4000 species, offers a plethora of cases for comparative and evolutionary studies. Despite a long history of phylogenetic inference, many relationships remain unresolved among the groups and genera in the Drosophilidae. To clarify these relationships, we first developed a set of new genomic markers and assembled a multilocus data set of 17 genes from 704 species of Drosophilidae. We then inferred well-supported group and species trees for this family. Additionally, we were able to determine the phylogenetic position of some previously unplaced species. These results establish a new framework for investigating the evolution of traits in fruit flies, as well as valuable resources for systematics.

## Introduction

The vinegar fly *Drosophila melanogaster* is a well-established and versatile model system in biology (Hales et al. 2015). The story began at the start of the 20^th^ century when the entomologist Charles Woodworth bred *D. melanogaster* in captivity, paving the way to seminal William Castle’s work at Harvard in 1901 (Sturtevant A. H. 1959). But it is undoubtedly with Thomas Hunt Morgan and his colleagues that *D. melanogaster* became a model organism in genetics (Morgan 1910). Nowadays, *D. melanogaster* research encompasses diverse fields, such as biomedicine (Ugur et al. 2016), developmental biology (Hales et al. 2015), growth control (Wartlick et al. 2011), gut microbiota (Trinder et al. 2017), innate immunity (Buchon et al. 2014), behaviour (Cobb 2007), and neuroscience (Bellen et al. 2010).

By the mid-20^th^ century, evolutionary biologists have widened *Drosophila* research by introducing many new species of Drosophilidae in comparative studies. For example, the mechanisms responsible for morphological differences of larval denticle trichomes (Sucena et al. 2003)(McGregor et al. 2007), adult pigmentation (Jeong et al. 2008)(Yassin, Delaney, et al. 2016), sex combs (Tanaka et al. 2009), and genital shape (Glassford et al. 2015)(Peluffo et al. 2015) have been thoroughly investigated across Drosophilidae. Comparative studies brought new insights into the evolution of ecological traits, such as host specialization (Lang et al. 2012)(Yassin et al. 2016), niche diversification (Chung et al. 2014), species distribution (Kellermann et al. 2009), pathogen virulence (Longdon et al. 2015), and behavior (Dai et al. 2008)(Karageorgi et al. 2017).

More than 150 genomes of *Drosophila* species are now sequenced (Adams et al. 2000)(Clark et al. 2007)(Wiegmann and Richards 2018)(Kim et al. 2020), allowing the comparative investigation of gene families (Sackton et al. 2007)(Almeida et al. 2014)(Finet et al. 2019) as well as global comparison of genome organization (Bosco et al. 2007)(Bhutkar et al. 2008). For all these studies, a clear understanding of the evolutionary relationships between species is necessary to interpret the results in an evolutionary context. A robust phylogeny is then crucial to confidently infer ancestral states, identify synapomorphic traits, and reconstruct the history of events during the evolution and diversification of Drosophilidae.

Fossil-based estimates suggest that the family Drosophilidae originated at least 30-50 Ma (Throckmorton 1975)(Grimaldi 1987)(Wiegmann et al. 2011). To date, the family comprises more than 4,392 species (DrosWLD-Species 2021) classified into two subfamilies, the Drosophilinae Rondani and the Steganinae Hendel. Each of these subfamilies contains several genera, which are traditionally subdivided into subgenera, and are further composed of species groups. Nevertheless, the monophyletic status of each of these taxonomic units is frequently controversial or unassessed. Part of this controversy is related to the frequent detection of paraphyletic taxa within Drosophilidae (Throckmorton 1975)(Katoh et al. 2000)(Robe et al. 2005)(Robe et al. 2010)(Da Lage et al. 2007)(Van Der Linde et al. 2010)(Russo et al. 2013)(Yassin 2013)(Katoh et al. 2017)(Gautério et al. 2020), although the absence of a consistent phylogenetic framework for the entire family makes it difficult to assess alternative scenarios.

Despite the emergence of the *Drosophila* genus as a model system to investigate the molecular genetics of functional evolution, relationships within the family Drosophilidae remain poorly supported. The first modern phylogenetic trees of this family relied on morphological characters (Throckmorton 1962)(Throckmorton 1975)(Throckmorton 1982), followed by a considerable number of molecular phylogenies that mainly focused on individual species groups (reviewed in (Markow and O’Grady 2006)(O’Grady and DeSalle 2018)). For the last decade, only a few large-scale studies have attempted to resolve the relationships within Drosophilidae as a whole. For example, supermatrix approaches brought new insights, such as the identification of the earliest branches in the subfamily Drosophilinae (Van Der Linde et al. 2010)(Yassin et al. 2010), the paraphyly of the subgenus *Drosophila* (*Sophophora*) (Gao et al. 2011), the placement of Hawaiian clades (O’Grady et al. 2011)(Lapoint et al. 2013)(Katoh et al. 2017), and the placement of Neotropical Drosophilidae (Lizandra J. Robe, Valente, et al. 2010). Most of the aforementioned studies have suffered from limited taxon or gene sampling. Recent studies improved the taxon sampling and the number of loci analysed (Morales-Hojas and Vieira 2012)(Russo et al. 2013)(Izumitani et al. 2016). To date, the most taxonomically-broad study is a revision of the Drosophilidae that includes 30 genera in Steganinae and 43 in Drosophilinae, but only considering a limited number of genomic markers (Yassin 2013).

To clarify the phylogenetic relationships in the Drosophilidae, we built a comprehensive dataset of 704 species that include representatives from most of the major genera, subgenera, and species groups in this family. We developed new genomic markers and compiled available ones from previously published phylogenetic studies. We then inferred well-supported trees at the group-and species-level for this family. Additionally, we were able to determine the phylogenetic position of several species of uncertain affinities. Our results establish a new framework for investigating the systematics and diversification of fruit flies and provide a valuable genomic resource for the *Drosophila* community.

## Results and Discussion

### A multigene phylogeny of 704 drosophilid species

We assembled a multilocus dataset of 17 genes (14,961 unambiguously aligned nucleotide positions) from 704 species of Drosophilidae. Our phylogeny recovers many of the clades or monophyletic groups previously described in the Drosophilidae (Figure 1). Whereas the branching of the species groups is mostly robust, some of the deepest branches of the phylogenic tree remain poorly supported or unresolved, especially in Bayesian analyses (see online supplementary tree files). This observation prompted us to apply a composite taxon strategy that has been used to resolve challenging phylogenetic relationships (Finet et al. 2010)(Campbell and Lapointe 2011)(Sigurdsen and Green 2011)(Charbonnier et al. 2015)(Mengual et al. 2017)(Fan et al. 2020). This approach limits branch lengths in selecting slow-evolving sequences, and decreases the percentage of missing data, allowing the use of parameter-rich models of evolution (Campbell and Lapointe 2009). We defined 63 composite groups as the monophyletic groups identified in the 704-taxon analysis (Figure 1, Table S1), and added these to the sequences of 20 other ungrouped taxa to perform additional phylogenetic evaluations. The overall bootstrap values and posterior probabilities were higher for the composite tree (Figures 2A, S1, and online supplementary tree files).

**Figure 1.**
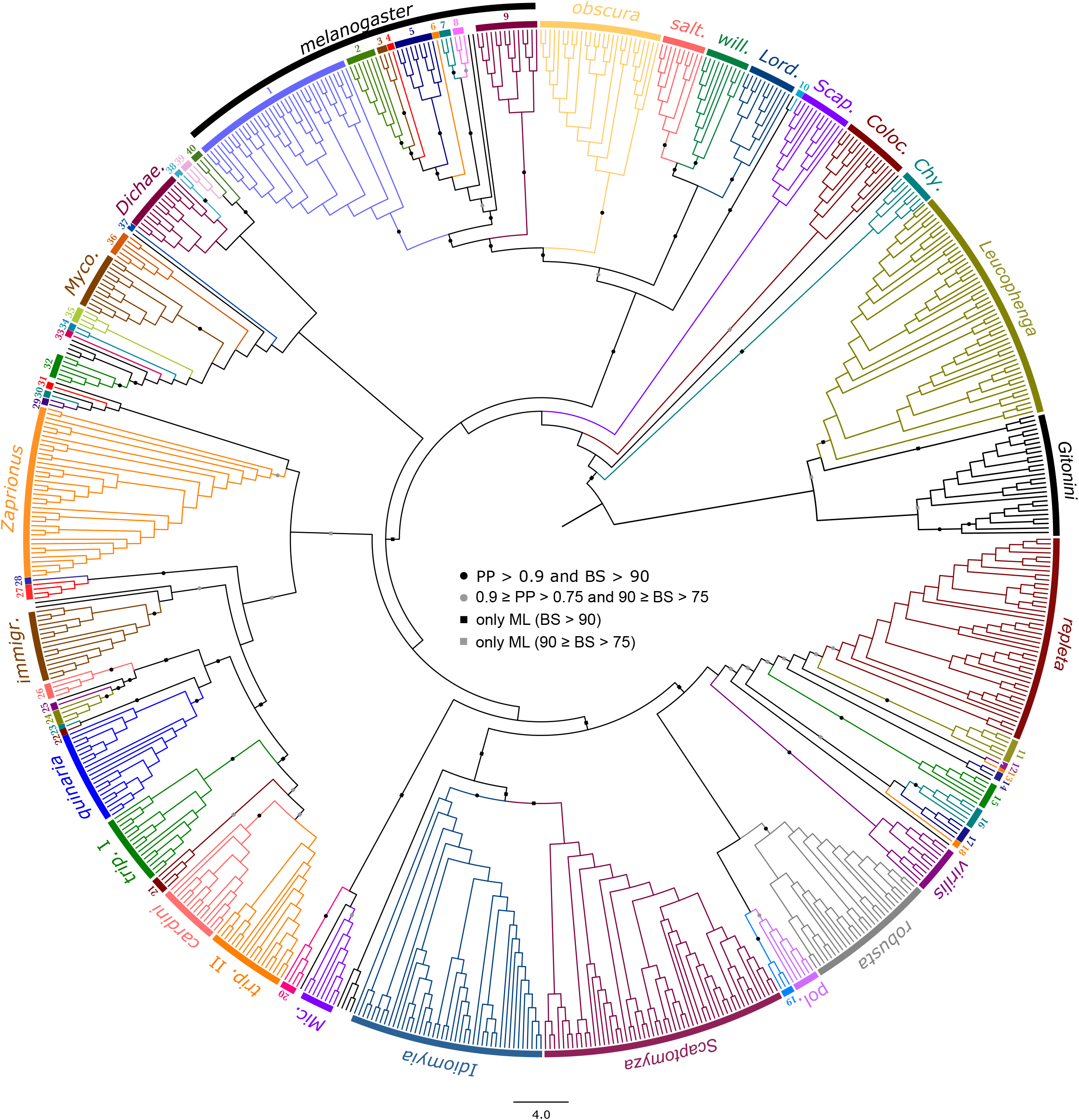
Phylogram of the 704-taxon analyses. IQ-TREE maximum-likelihood analysis was conducted under the GTR+R+FO model. Support values obtained after 100 bootstrap replicates are shown for selected supra-group branches, and infra-group branches within the *melanogaster* group (all the support values are shown online). Black dots indicate support values of PP > 0.9 and BP > 90; grey dots 0.9 ≥ PP > 0.75 and 90 ≥ BP > 75; black squares only BP > 90; grey squares only 90 ≥ BP > 75. Scale bar indicates the number of changes per site. Groups and subgroups are numbered or abbreviated as follows: (1) *montium*, (2) *takahashii* sgr, (3) *suzukii* sgr, (4) *eugracilis* sgr, (5) *melanogaster* sgr, (6) *ficusphila* sgr, (7) *elegans* sgr, (8) *rhopaloa* sgr, (9) *ananassae*, (10) *Collessia*, (11) *mesophragmatica*, (12) *dreyfusi*, (13), *coffeata*, (14) *canalinea*, (15) *nannoptera*, (16) *annulimana*, (17) *flavopilosa*, (18) *flexa*, (19) *angor*, (20) *Dorsilopha*, (21) *ornatifrons*, (22) *histrio*, (23) *macroptera*, (24) *testacea*, (25) *bizonata*, (26) *funebris*, (27) *Samoaia*, (28) *quadrilineata* sgr, (29) *Liodrosophila*, (30) *Hypselothyrea*, (31) *Sphaerogastrella*, (32) *Zygothrica* I, (33) *Paramycodrosophila*, (34) *Hirtodrosophila* III, (35) *Hirtodrosophila* II, (36) *Hirtodrosophila* I, (37) *Dettopsomyia*, (38) *Mulgravea*, (39) *Hirtodrosophila* IV, (40) *Zygothrica* II, *Chy*: *Chymomyza*; *Colo*: *Colocasiomyia*; *Dichae*: *Dichaetophora*; *immigr*: *immigrans*; *Lord*: *Lordiphosa*; *Mic*: *Microdrosophila*; *Myco*: *Mycodrosophila*; *pol*: *polychaeta*; *salt*: *saltans*; *Scap*: *Scaptodrosophila*; *trip*: *tripunctata*; *will*: *willistoni*.

**Figure 2.**
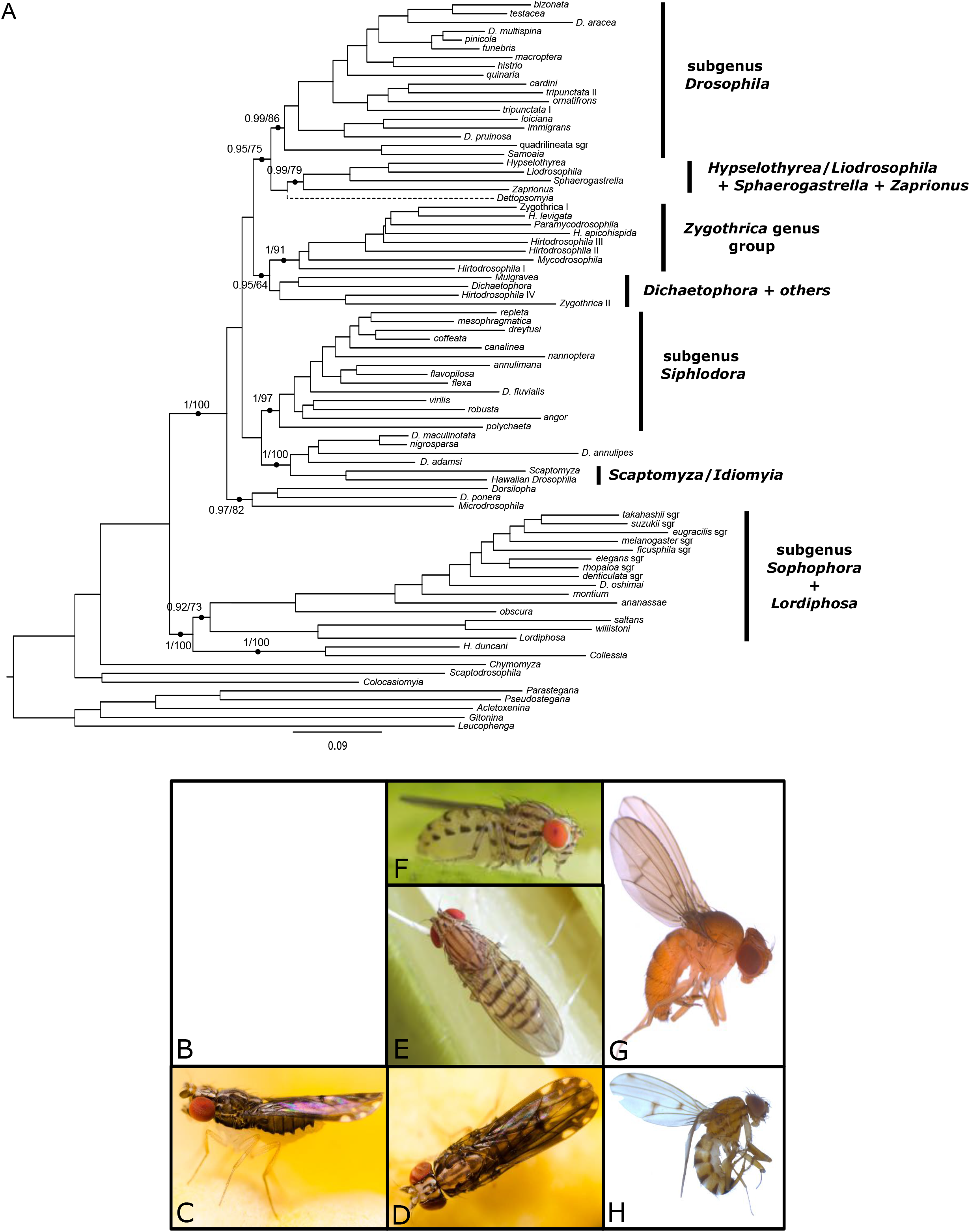
(A) Phylogram of the 83-taxon analyses. The overall matrix represents 14,961 nucleotides and 83 taxa, including 63 composite ones. Support values obtained after 100 bootstrap replicates and Bayesian posterior probabilities are shown for selected branches and mapped onto the ML topology (all the support values are shown in Figure S1). The dotted line indicates that the placement of *Dettopsomyia* varies between ML and Bayesian trees. Scale bar indicates the number of changes per site. (B-H) Photos of species of particular interest in this paper. (B) *Drosophila oshimai* female (top) and male (bottom) (Japan, courtesy of Japan Drosophila Database), (C-D) *Collessia kirishimana* (Japan, courtesy of Masafumi Inoue), (E-F) *Drosophila annulipes* (Japan, courtesy of Yasuo Hoshino), (G) *Drosophila pruinosa* (São Tomé, courtesy of Stéphane Prigent), (H) *Drosophila adamsi* (Cameroun, courtesy of Stéphane Prigent).

Incongruence among phylogenetic markers is a common source of error in phylogenomics (Jeffroy et al. 2006). In order to estimate the presence of incongruent signal in our dataset, we first investigated the qualitative effect of single marker removal on the topology of the composite tree (Figure S2). We found the overall topology is very robust to marker sampling, with only a few minor changes for each dataset. For instance, the *melanogaster* subgroup sometimes clusters with the *eugracilis* subgroup instead of branching off prior to the *eugracilis* subgroup (Figures 2 and S2). The position of the genus *Dettopsomyia* and that of the *angor* and *histrio* groups is also very sensitive to single marker removal, which could explain the low support values obtained (Figures 2 and S2). To a lesser extent, the position of *D. fluvialis* can vary as well depending on the removed marker (Figures 2 and S2). We also quantitatively investigated the incongruence present in our dataset by calculating genealogical concordance. The gene concordance factor is defined as the percentage of individual gene trees containing that node for every node of the reference tree. Similarly, the fraction of nodes supported by each marker can be determined. The markers we developed in this study show concordance rates ranging from 46.2 to 90.9% (Figure 3, Table 2). With an average concordance rate of 65%, these new markers appear as credible phylogenetic markers, without significantly improving the previous markers (average concordance rate of 64.8%).

**Table 1.**
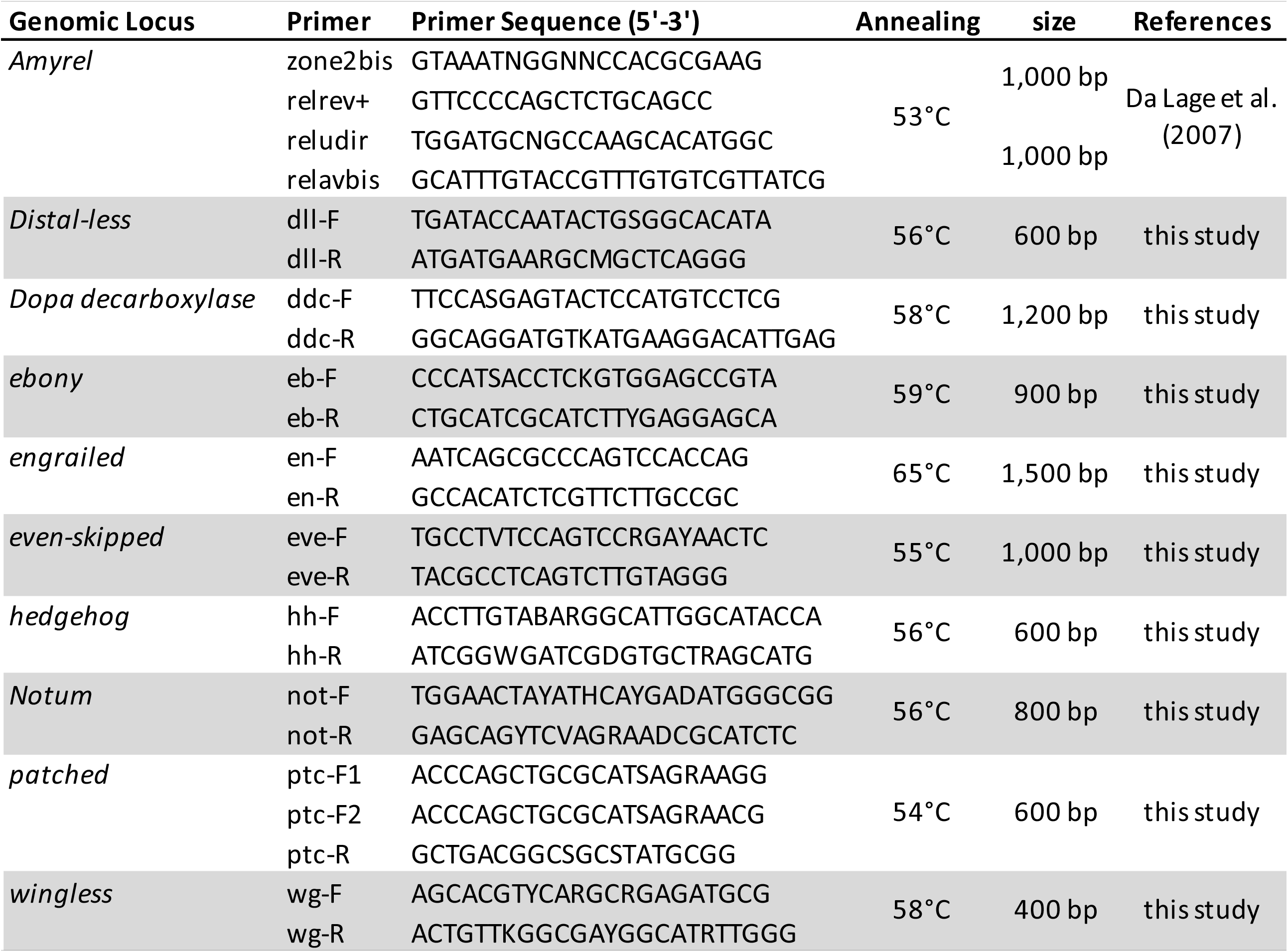
List of PCR primers used in this study.

**Table 2.**
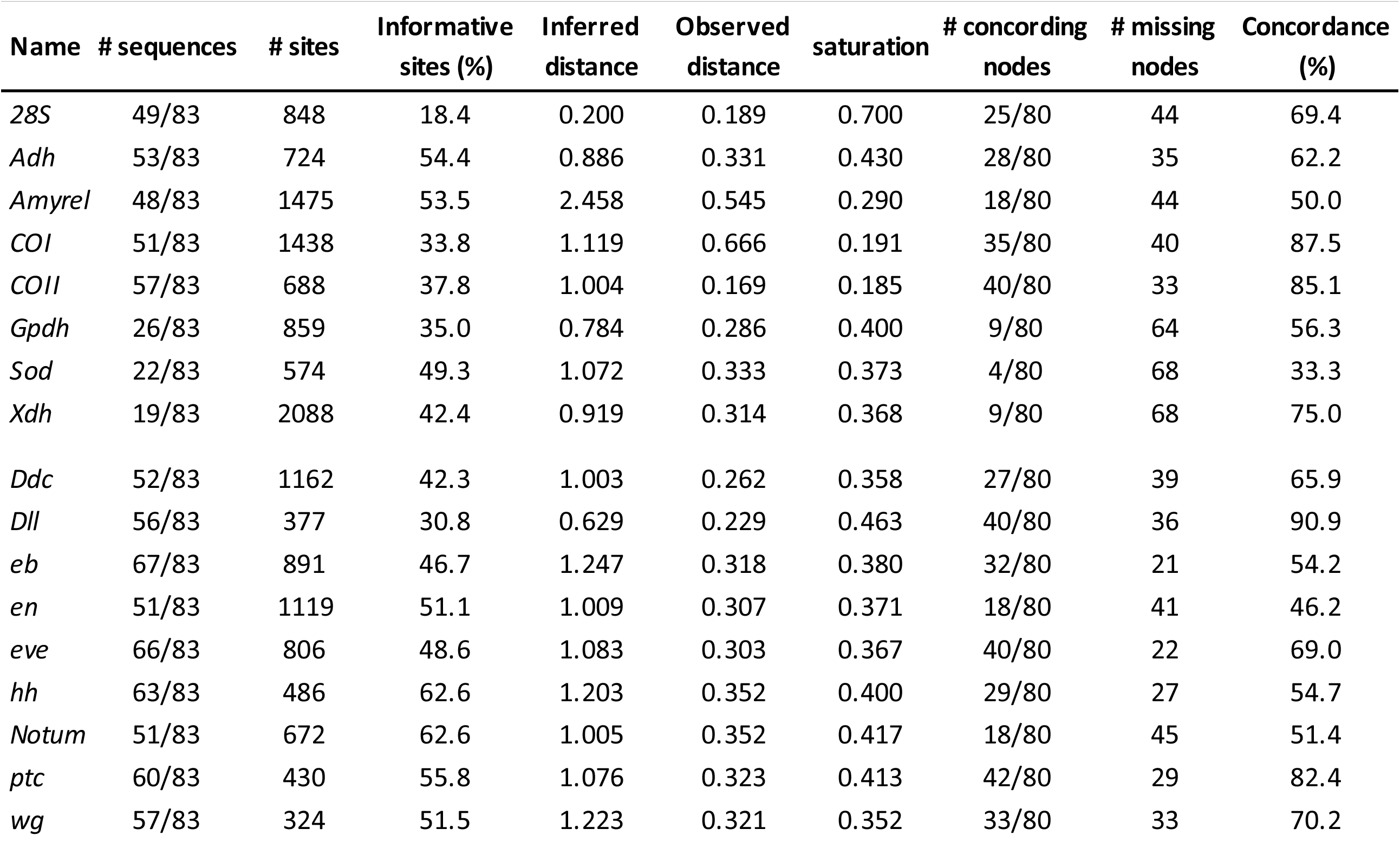
Dataset statistics.

**Figure 3.**
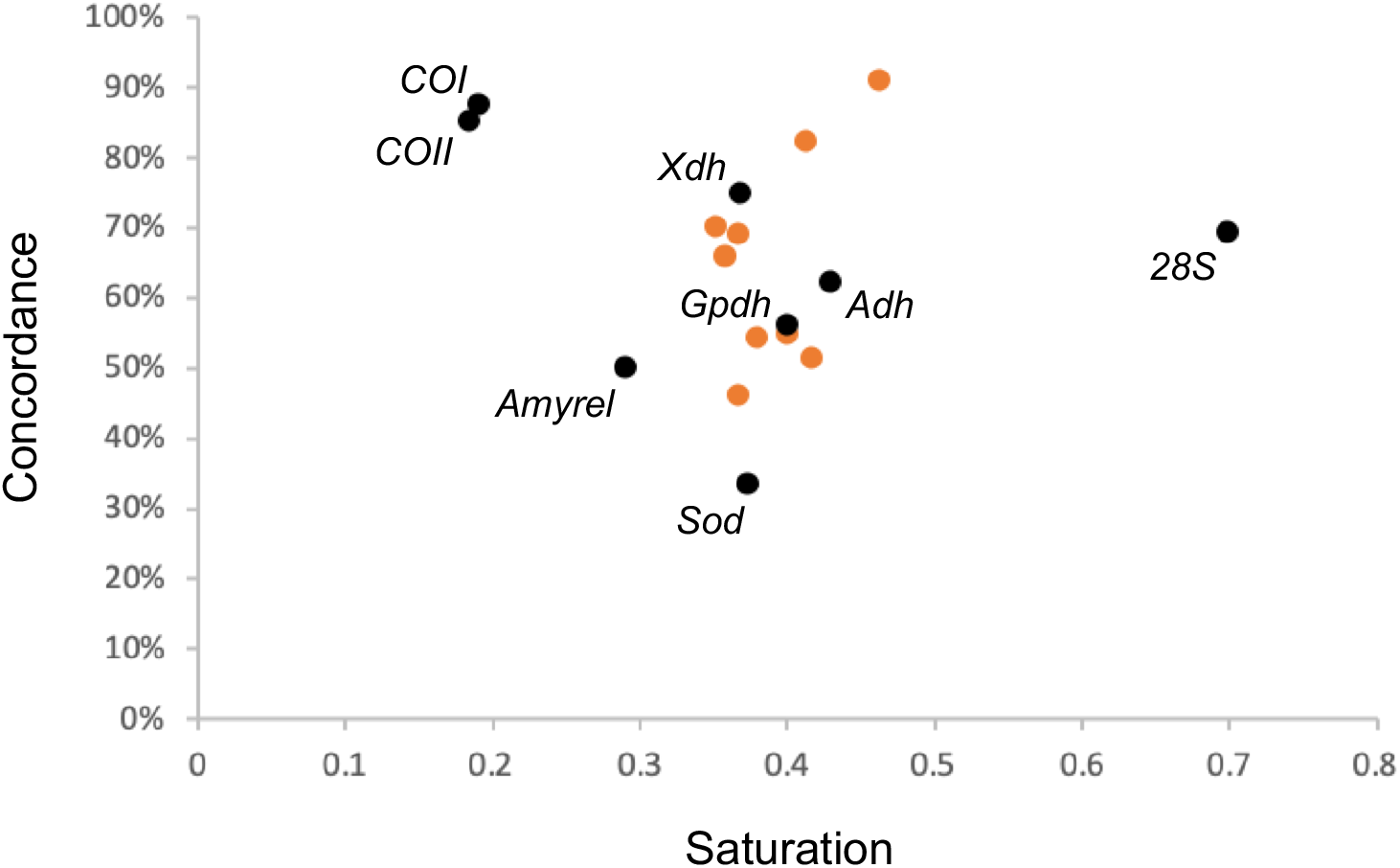
Concordance *versus* mutational saturation of the phylogenetic markers. The y-axis indicates the percentage of concordant nodes, and the x-axis indicates the saturation level. In comparison with published markers (black dots), the markers developed in this study (orange dots) generally show moderate saturation levels and satisfying concordance.

Multiple substitutions at the same position is another classical bias in phylogenetic reconstruction, capable of obscuring the genuine phylogenetic signal (Jeffroy et al. 2006). We quantified the mutational saturation for each phylogenetic marker. On average, the newly developed markers are moderately saturated (Figure 3, Table 2). These markers are indeed less saturated than the *Amyrel, COI*, and *COII* genes that have been commonly applied for phylogenetic inference in Drosophilidae (Baker and Desalle 1997)(O’Grady et al. 1998)(Remsen and O’Grady 2002)(Bonacum et al. 2005)(Da Lage et al. 2007)(Robe et al. 2010)(Gao et al. 2011)(O’Grady et al. 2011)(Russo et al. 2013)(Yassin 2013).

In the following sections of the paper, we will highlight and discuss some of the most interesting results we obtained. Our analyses either confirm or challenge previous phylogenies, and shed light on several unassessed questions, contributing to an emerging picture of phylogenetic relationships in Drosophilidae.

### The *Sophophora* subgenus and closely related taxa

We found that the *obscura*-*melanogaster* clade is the sister group of the lineages formed by the Neotropical *saltans* and *willistoni* groups, and the *Lordiphosa* genus (Bayesian posterior probability [PP] = 0.92, bootstrap percentage [BP] = 73) (Figures 2A and S1). Thus, our study recovers the relationship between the groups of the *Sophophora* subgenus (Gao et al. 2011)(Russo et al. 2013)(Yassin 2013) and supports the paraphyletic status of *Sophophora* regarding *Lordiphosa* (Katoh et al. 2000). However, we noted substantial changes within the topology presented for the *melanogaster* species group. The original description of *Drosophila oshimai* noted a likeness to *Drosophila unipectinata*, thus classifying *D. oshimai* into the *suzukii* species subgroup (Choo and Nakamura 1973). The phylogenetic tree we obtained does not support this classification (Figure 2A). It rather defines *D. oshimai* as the representative of a new subgroup (PP = 1, BP = 96) that diverged immediately after the split of the *montium* group. The position of *D. oshimai* therefore challenges the monophyly of the *suzukii* subgroup. Interestingly, the paraphyly of the *suzukii* subgroup has also been suggested in previous studies (Lewis et al. 2005)(Russo et al. 2013). Another interesting case is the positioning of the *denticulata* subgroup that has never been tested before. Our analysis convincingly places its representative species *Drosophila denticulata* as the fourth subgroup to branch off within the *melanogaster* group (PP = 1, BP = 82). Last, the topology within the *montium* group drastically differs from the most recent published phylogeny (Conner et al. 2021).

The genus *Collessia* comprises five described species that can be found in Australia, Japan, and Sri Lanka, but its phylogenetic status was so far quite ambiguous (Okada 1967)(Bock 1982)(Okada 1988). In addition, Grimaldi (1990) proposed that *Tambourella ornata* should belong to the genus *Collessia*. These two genera are similar in the wing venation and pigmentation pattern (Okada 1984).

Our phylogenetic analysis identifies *Collessia* as sister group to the species *Hirtodrosophila duncani* (PP = 1, BP = 100). Interestingly, this branching is also supported by morphological similarities shared between the genera *Collessia* and *Hirtodrosophila*. The species *C. kirishimana* and *C. hiharai* were indeed initially described as *Hirtodrosophila* species (Okada 1967) before being assigned to the genus *Collessia* (Okada 1984). The clade *Collessia*-*H. duncani* is sister to the *Sophophora*-*Lordiphosa* lineage in the ML inference (BP = 100) but to the Neotropical *Sophophora*-*Lordiphosa* clade in the Bayesian inference (PP = 0.92).

### The early lineage of *Microdrosophila* and *Dorsilopha*

Within the tribe Drosophilini, all the remaining taxa (composite taxa + ungrouped species) other than those of the *Sophophora*-*Lordiphosa* and *Collessia-H. duncani* lineage form a large clade (PP = 1, BP = 100). Within this clade, the genus *Microdrosophila*, the subgenus *Dorsilopha*, and *Drosophila ponera* group into a lineage (PP = 0.97, BP = 82) that appears as an early offshoot (PP = 1.00, BP = 59). *Drosophila ponera* is an enigmatic species collected in La Réunion (David and Tsacas 1975), whose phylogenetic position has never or rarely been investigated. In spite of morphological similarities with the *quinaria* group, the authors suggested to keep *D. ponera* as ungrouped with respect to a divergent number of respiratory egg filaments (David and Tsacas 1975). To our knowledge, our study is the first attempt to phylogenetically position this species. We found that *D. ponera* groups with the *Dorsilopha* subgenus (PP = 0.99, BP = 75) within this early-diverging lineage.

### The Hawaiian drosophilid clade and the *Siphlodora* subgenus

The endemic Hawaiian Drosophilidae contain approximately 1,000 species that split into the Hawaiian *Drosophila* (or *Idiomyia* genus according to Grimaldi (1990)) and the genus *Scaptomyza* (O’Grady et al. 2009). Generally considered as sister to the *Siphlodora* subgenus (Robe et al. 2010)(Russo et al. 2013)(Yassin 2013), these lineages represent a remarkable framework to investigate evolutionary radiation and subsequent diversification of morphology (Stark and O’Grady 2010), pigmentation (Edwards et al. 2007), ecology (Magnacca et al. 2008), and behavior (Kaneshiro 1999). Although the relationships within the *Siphlodora* clade are generally in agreement with previous studies (Tatarenkov et al. 2001)(Robe et al. 2010)(Russo et al. 2013)(Yassin 2013), its sister clade does not seem to be restricted to the Hawaiian Drosophilidae. In fact, according to our phylogenies, it also includes at least four other species of the genus *Drosophila* (Figures 2A, S1, and online supplementary tree files). We propose that this broader clade, rather than the Hawaiian clade *sensu stricto*, should be seen as a major lineage of Drosophilidae.

This broader clade is strongly supported (PP = 1, BP = 100) and divided into two subclades, one comprises the genera *Idiomyia* and *Scaptomyza* (PP = 0.99, BP = 97) and the other includes *D. annulipes, D. adamsi, D. maculinotata* and *D. nigrosparsa* (PP = 0.99, BP = 75). The latter subclade, also suggested by Katoh et al. (2007) and Russo et al. (2013), is interesting with respect to the origin of Hawaiian drosophilids. Of the four component species, *D. annulipes* was originally described as a member of the subgenus *Spinulophila*, which was synonymized with *Drosophila* and currently corresponds to the *immigrans* group, although Wakahama et al. (1983) and Zhang and Toda (1992) cast doubt on its systematic position. As for *D. adamsi*, Da Lage et al. (2007) suggested it may be close to the *Idiomyia*-*Scaptomyza* clade, which is supported by our analyses. On the other hand, Prigent et al. (2013) based on morphological characters and Prigent et al. (2017) based on DNA barcoding have proposed that *D. adamsi* defines a new species group along with *D. acanthomera* and an undescribed species. *Drosophila adamsi* resembles *D. annulipes* in the body color pattern (Fig. 2F,E,H), suggesting their close relationship: Adams (1905) described, “mesonotum with five longitudinal, brown vittae, the central one broader than the others and divided longitudinally by a hair-like line, …; scutellum yellow, with two sublateral, brownish lines, …; pleurae with three longitudinal brownish lines”, for *Drosophila quadrimaculata* Adams, 1905, which is a homonym of *Drosophila quadrimaculata* Walker, 1856 and has been replaced with the new specific epithet “*adamsi*” by Wheeler (1959). Another species, *D. nigrosparsa*, belongs to the *nigrosparsa* species group, along with *D. secunda, D. subarctica* and *D. vireni* (Bächli et al. 2004). Moreover, Máca (1992) pointed out the close relatedness of *D. maculinotata* to the *nigrosparsa* group.

### The *Drosophila* subgenus and closely related taxa

Although general relationships within the *Drosophila* subgenus closely resemble those recovered by previous studies (Hatadani et al. 2009)(Robe et al. 2010)(Robe et al. 2010)(Izumitani et al. 2016), there are some outstanding results related to other genera or poorly studied *Drosophila* species.

*Samoaia* is a small genus of seven described species endemic to the Samoan Archipelago (Malloch 1934)(Wheeler and Kambysellis 1966), particularly studied for their body and wing pigmentation (Dufour et al. 2020). In our analysis, the genus *Samoaia* is found to group with the *quadrilineata* species subgroup of the *immigrans* group. This result is similar to conclusions formulated by some previous studies (Tatarenkov et al. 2001)(Robe et al. 2010)(Yassin et al. 2010)(Yassin 2013), but differs from other published phylogenies in which *Samoaia* is sister to most other lineages in the subgenus *Drosophila* (Russo et al. 2013). It is noteworthy that our sampling is the most substantial with four species of *Samoaia*.

The two African species *Drosophila pruinosa* and *Drosophila pachneissa*, which were assigned to the *loiciana* species complex because of shared characters such as a glaucous-silvery frons and rod-shaped surstyles (Tsacas 2002), are placed together with the *immigrans* group (PP = 1, BP = 94). In previous large-scale analyses, *D. pruinosa* was suggested to group with *Drosophila sternopleuralis* into the sister clade of the *immigrans* group (Da Lage et al. 2007)(Russo et al. 2013).

Among other controversial issues, the phylogenetic position of *Drosophila aracea* was previously found to markedly change according to the phylogenetic reconstruction methods (Da Lage et al. 2007). This anthophilic species lives in Central America (Heed and Wheeler 1957). Its name comes from the behavior of females that lay eggs on the spadix of plants in the family Araceae (Heed and Wheeler 1957)(Tsacas and Chassagnard 1992). Our analysis places *D. aracea* as the sister taxon of the *bizonata*-*testacea* clade with high confidence (PP = 1, BP = 85). No occurrence of flower-breeding behavior has been reported in the *bizonata*-*testacea* clade, reinforcing the idea that *D. aracea* might have recently evolved from a generalist ancestor (Tsacas and Chassagnard 1992).

### The *Zygothrica* genus group

The fungus-associated genera *Hirtodrosophila, Mycodrosophila, Paraliodrosophila, Paramycodrosophila*, and *Zygothrica* contain 448 identified species (TaxoDros 2020) and have been associated with the *Zygothrica* genus group (Grimaldi 1990). Although the *Zygothrica* genus group was recurrently recovered as paraphyletic (Da Lage et al. 2007)(Van Der Linde et al. 2010)(Russo et al. 2013)(Yassin 2013), two recent studies suggest, on the contrary, its monophyly (Gautério et al. 2020)(Zhang et al. 2021). Our study does not support the monophyly of the *Zygothrica* genus group in virtue of the polyphyletic status of *Hirtodrosophila* and *Zygothrica*: some representatives (*e*.*g*., *H. duncani)* cluster with *Collessia*, while others (*e*.*g*., *Hirtodrosophila* IV and *Zygothrica* II) appear closely related to the genera *Dichaetophora* and *Mulgravea*. Furthermore, the placement of the *Zygothrica* genus group recovered in our study also differs from some previous estimates. In fact, the broadly defined *Zygothrica* genus group, which includes *Dichaetophora* and *Mulgravea* (PP = 0.95, BP = 64), appears as sister to the clade composed of the subgenus *Drosophila* and the *Hypselothyrea*/*Liodrosophila* + *Sphaerogastrella* + *Zaprionus* clade (PP = 1, BP = 56) (Figures 2A and S1). This placement is similar to the ones obtained in different studies (Van Der Linde et al. 2010)(Russo et al. 2013), but contrasts with the close relationship of the *Zygothrica* genus group to the subgenus *Siphlodora* + *Idiomyia*/*Scaptomyza* proposed in two recent studies (Gautério et al. 2020)(Zhang et al. 2021). Given the moderate bootstrap value, the exact status of the *Zygothrica* genus group remains as an open question. Furthermore, within the superclade of the broadly defined *Zygothrica* genus group (Figures 1 and 2A), the genus *Hirtodrosophila* is paraphyletic and split into four independent lineages, reinforcing previous suggestions based on multilocus approaches (Van Der Linde et al. 2010)(Gautério et al. 2020)(Zhang et al. 2021). This also occurred with the genus *Zygothrica*, which split into two independent clades (Figure 2A). The *leptorostra* subgroup (*Zygothrica* II) clusters with the subgroup *Hirtodrosophila* IV (PP = 1, BP = 100), whereas the *Zygothrica* I subgroup clusters with the species *Hirtodrosophila levigata* (PP = 0.99, BP = 98).

### DrosoPhyla: a powerful tool for systematics

Besides bringing an updated and improved phylogenetic framework to Drosophilidae, our approach also addresses several questions that were previously unassessed or controversial at the genus, subgenus, group, or species level. We are therefore confident that it may become a powerful tool for future drosophilid systematics. According to diversity surveys (O’Grady and DeSalle 2018), ∼25% of drosophilid species remain to be discovered, potentially a thousand species to place in the tree of Drosophilidae. While whole-genome sequencing is becoming widespread, newly discovered species often come down to a few specimens pinned or stored in ethanol – non-optimal conditions for subsequent genome sequencing and whole-genome studies. Based on a few short genomic markers, our approach is compatible with taxonomic work, and gives good resolution.

## Acknowledgements

We thank Jean-Luc Da Lage and John Jaenike for providing fly specimens. We thank Virginie Orgogozo and Noah Whiteman for giving early access to the genome of *D. pachea* and *S. flava*, respectively. We thank Masafumi Inoue, Stéphane Prigent, Yasuo Hoshino, and the Japan Drosophila Database for providing photos. We thank Amir Yassin for fruitful discussions and comments on the manuscript. We thank the Sean Carroll laboratory for discussions and financial support.

## Material and Methods

### Taxon sampling

The species used in this study were sampled from different locations throughout the world (Table S1). The specimens were field-collected by the authors, purchased from the National Drosophila Species Stock Center (http://blogs.cornell.edu/drosophila/) and the Kyoto Stock Center (https://kyotofly.kit.jp/cgi-bin/stocks/index.cgi), or obtained from colleagues. Individual flies were preserved in 100% ethanol and identified based on morphological characters.

### Data collection

Ten genomic markers were amplified by PCR using degenerate primers developed for the present study (Table 1). Genomic DNA was extracted from a single adult fly as follows: the fly was placed in a 0.5-mL tube and mashed in 50 μL of squishing buffer (Tris-HCl pH=8.2 10 mM, EDTA 1 mM, NaCl 25 mM, proteinase K 200 μg/mL) for 20-30 seconds, the mix was incubated at 37°C for 30 minutes, then the proteinase K was inactivated by heating at 95°C for 1-2 minutes. A volume of 1 μL was used as template for PCR amplification. Nucleotide sequences were also retrieved from the NCBI database for the five nuclear markers *28S ribosomal RNA* (*28S*), *alcohol dehydrogenase* (*Adh*), *glycerol-3-phosphate dehydrogenase* (*Gpdh*), *superoxide dismutase* (*Sod*), *xanthine dehydrogenase* (*Xdh*), and the two mitochondrial markers *cytochrome oxidase subunit 1* (*COI*) and *cytochrome oxidase subunit 2* (*COII*). The sequences reported in this paper have been deposited in GenBank under specific accession numbers: *Amyrel* (MW392482-MW392524), *Ddc* (MW403139-MW403307), *Dll* (MW403308-MW403483), *eb* (MW415022-MW415267), *en* (MW418945-MW419079), *eve* (MW425034-MW425273), *hh* (MW385549-MW385782), *Notum* (MW429853-MW430003), *ptc* (MW442160-MW442361), *wg* (MW392301-MW392481).

### Phylogenetic reconstruction

Alignments for each individual gene were generated using MAFFT 7.45 (Katoh and Standley 2013), and unreliably aligned positions were excluded using trimAl with parameters -gt 0.5 and -st 0.001 (Capella-Gutiérrez et al. 2009). The possible contamination status was verified by inferring independent trees for each gene using RAxML 8.2.4 under the GTR+Г model (Stamatakis 2014). Thus, any sequence leading to the suspicious placement of a taxonomically well-assigned species was removed from the dataset. Moreover, almost identical sequences leading to very short tree branches were carefully examined and excluded if involving non-closely related taxa. In-house Python scripts (available on GitHub XXX) were used to concatenate the aligned and filtered sequences, and the resulting dataset was used for phylogenetic reconstruction. Maximum-likelihood (ML) searches were performed using IQ-TREE 2.0.6 (Minh, Schmidt, et al. 2020) under the GTR model, with the FreeRate model of rate heterogeneity across sites with four categories, and ML estimation of base frequencies from the data (GTR+R+FO). The edge-linked proportional partition model was used with one partition for each gene. Sequence alignments and tree files are available from (https://www.dropbox.com/sh/ts2pffqnnwd34c8/AAA9qLL7dCC3urxR1NcioJvLa?dl=0).

### Composite taxa

This strategy started from clustering the species by unambiguous monophyletic genera, groups, or subgroups identified in the 704-taxon analysis. After this, the least diverging sequence or species recovered for each taxonomic unit for each marker was selected to ultimately yield a unique composite taxon by concatenation. The composite matrix was also used for conducting ML and Bayesian phylogenetic inference using IQ-TREE under a partitioned GTR+R+FO model, and PhyloBayes under a GTR+Г model (Lartillot et al. 2009), respectively. Sequence alignments and tree files are available from XXX.

### Saturation and concordance analysis

For each marker gene, the saturation was computed by performing a simple linear regression of the percent identity for each pair of taxa (observed distance) onto the ML patristic distance (inferred distance) (Philippe et al. 1994) estimated using the ETE 3 library (Huerta-Cepas et al. 2016). We also calculated per gene and per site concordance factors using IQ-TREE under the GTR+R+FO model as recently described (Minh, Hahn, et al. 2020).

## Figure legends

**Figure S1**. Phylogram of the 83-taxon analyses. (Left) IQ-TREE maximum-likelihood analyses were conducted using the GTR+R+FO model. Support values obtained after 100 bootstrap replicates are shown for all branches. Scale bar indicates the number of changes per site. (Right) PhyloBayes Bayesian analyses were conducted using the GTR+Г model. Bayesian posterior probabilities are shown for all branches. Scale bar indicates the number of changes per site.

**Figure S2**. The impact of marker sampling on the tree topology. The composite tree was built on 17 different datasets that correspond to the whole dataset minus one marker sequentially removed. The changes in relation to the ML composite tree depicted in Figure 2 are shown in red. Scale bar indicates the number of changes per site.

**Figure S3**. Mutational saturation of the 17 phylogenetic markers. The x-axis indicates 799 the distance inferred from the ML composite tree, whereas the y-axis indicates the 800 observed distance between two taxa. The slope of the red line is an indicator of the 801 saturation level, low values meaning high saturation. The black line corresponds to 802 the absence of multiple substitutions.

## Table legends

**Table S1**. Taxon sampling.

## References

Adams CF. 1905. Diptera Africana, I. Kansas Univ. Sci. Bull. 3:149–188.

Adams MD, Celniker SE, Holt RA, Evans CA, Gocayne JD, Amanatides PG, Scherer SE, Li PW, Hoskins RA, Galle RF, et al. 2000. The genome sequence of Drosophila melanogaster. Science 287:2185–2195.

Almeida FC, Sánchez-Gracia A, Campos JL, Rozas J. 2014. Family size evolution in drosophila chemosensory gene families: A comparative analysis with a critical appraisal of methods. Genome Biol. Evol. 6:1669–1682.

Baker RH, Desalle R. 1997. Multiple sources of character information and the phylogeny of Hawaiian Drosophilids. Syst. Biol. 46:654–673.

Bellen HJ, Tong C, Tsuda H. 2010. 100 years of Drosophila research and its impact on vertebrate neuroscience: a history lesson for the future. Nat. Rev. Neurosci. 11:514–522.

Bhutkar A, Schaeffer SW, Russo SM, Xu M, Smith TF, Gelbart WM. 2008. Chromosomal rearrangement inferred from comparisons of 12 drosophila genomes. Genetics 179:1657–1680.

Bock I. 1982. Drosophilidae of Australia V. Remaining genera and synopsis (Insecta: Diptera). Aust. J. Zool. 89:1–164.

Bonacum J, O’Grady PM, Kambysellis M, DeSalle R. 2005. Phylogeny and age of diversification of the planitibia species group of the Hawaiian Drosophila. Mol. Phylogenet. Evol. 37:73–82.

Bosco G, Campbell P, Leiva-Neto JT, Markow TA. 2007. Analysis of Drosophila species genome size and satellite DNA content reveals significant differences among strains as well as between species. Genetics 177:1277–1290.

Buchon N, Silverman N, Cherry S. 2014. Immunity in Drosophila melanogaster — from microbial recognition to whole-organism physiology. Nat. Rev. Immunol. 14:796–810.

Campbell V, Lapointe FJ. 2009. The use and validity of composite taxa in phylogenetic analysis. Syst. Biol. 58:560–572.

Campbell V, Lapointe FJ. 2011. Retrieving a mitogenomic mammal tree using composite taxa. Mol. Phylogenet. Evol. 58:149–156.

Capella-Gutiérrez S, Silla-Martínez JM, Gabaldón T. 2009. trimAl: A tool for automated alignment trimming in large-scale phylogenetic analyses. Bioinformatics 25:1972–1973.

Charbonnier S, Audo D, Barriel V, Garassino A, Schweigert G, Simpson M. 2015. Phylogeny of fossil and extant glypheid and litogastrid lobsters (Crustacea, Decapoda) as revealed by morphological characters. Cladistics 31:231–249.

Choo J, Nakamura K. 1973. On a new species of Drosophila (Sophophora) from Japan (Diptera). Kontyû 41:305–306.

Chung H, Loehlin DW, Dufour HD, Vaccarro K, Millar JG, Carroll SB. 2014. A single gene affects both ecological divergence and mate choice in Drosophila. Science 343:1148–1151.

Clark AG, Eisen MB, Smith DR, Bergman CM, Oliver B, Markow TA, Kaufman TC, Kellis M, Gelbart W, Iyer VN, et al. 2007. Evolution of genes and genomes on the Drosophila phylogeny. Nature.

Cobb M. 2007. A gene mutation which changed animal behaviour: Margaret Bastock and the yellow fly. Anim. Behav. 74:163–169.

Conner WR, Delaney EK, Bronski MJ, Ginsberg PS, Wheeler TB, Richardson KM, Peckenpaugh B, Kim KJ, Watada M, Hoffmann AA, et al. 2021. A phylogeny for the Drosophila montium species group: A model clade for comparative analyses. Mol. Phylogenet. Evol. 158:107061.

Dai H, Chen Y, Chen S, Mao Q, Kennedy D, Landback P, Eyre-Walker A, Du W, Long M. 2008. The evolution of courtship behaviors through the origination of a new gene in Drosophila. Proc. Natl. Acad. Sci. U. S. A. 105:7478–7483.

David J, Tsacas L. 1975. Les Drosophilidae (Diptera) de l’Ile de la Réunion et de l’Ile Maurice. I. Deux nouvelles espèces du genre Drosophila. Bull. Mens. la Société Linnéenne Lyon 5:134–143.

DrosWLD-Species. 2021. DrosWLD-Species.

Dufour HD, Koshikawa S, Finet C. 2020. Temporal flexibility of gene regulatory network underlies a novel wing pattern in flies. Proc. Natl. Acad. Sci. 117:11589–11596.

Edwards KA, Doescher LT, Kaneshiro KY, Yamamoto D. 2007. A database of wing diversity in the Hawaiian Drosophila. PLoS One 2:3487.

Fan L, Wu D, Goremykin V, Xiao J, Xu Y, Garg S, Zhang C, Martin WF, Zhu R. 2020. Phylogenetic analyses with systematic taxon sampling show that mitochondria branch within Alphaproteobacteria. Nat. Ecol. Evol. 4:1213–1219.

Finet C, Slavik K, Pu J, Carroll SB, Chung H. 2019. Birth-and-Death Evolution of the Fatty Acyl-CoA Reductase (FAR) Gene Family and Diversification of Cuticular Hydrocarbon Synthesis in Drosophila. Genome Biol. Evol. 11:1541–1551.

Finet C, Timme RE, Delwiche CF, Marlétaz F. 2010. Multigene phylogeny of the green lineage reveals the origin and diversification of land plants. Curr. Biol. 20:2217–2222.

Gao JJ, Hu YG, Toda MJ, Katoh T, Tamura K. 2011. Phylogenetic relationships between Sophophora and Lordiphosa, with proposition of a hypothesis on the vicariant divergences of tropical lineages between the Old and New Worlds in the family Drosophilidae. Mol. Phylogenet. Evol. 60:98–107.

Gautério TB, Machado S, Loreto EL da S, Gottschalk MS, Robe LJ. 2020. Phylogenetic relationships between fungus-associated Neotropical species of the genera Hirtodrosophila, Mycodrosophila and Zygothrica (Diptera, Drosophilidae), with insights into the evolution of breeding sites usage. Mol. Phylogenet. Evol. 145.

Glassford WJ, Johnson WC, Dall NR, Smith SJ, Liu Y, Boll W, Noll M, Rebeiz M. 2015. Co-option of an Ancestral Hox-Regulated Network Underlies a Recently Evolved Morphological Novelty. Dev. Cell 34:520–531.

Grimaldi D. 1987. Amber Fossil Drosophilidae (Diptera), with Particular Reference to the Hispaniolan taxa. Am. Museum Novit. 2880:1–23.

Grimaldi DA. 1990. A Phylogenetic, Revised Classification of Genera in the Drosophilidae (Diptera). Bull. Am. Museum Nat. Hist. 197.

Hales KG, Korey CA, Larracuente AM, Roberts DM. 2015. Genetics on the fly: A primer on the drosophila model system. Genetics 201:815–842.

Hatadani LM, McInerney JO, Medeiros HF de, Junqueira ACM, Azeredo-Espin AM de, Klaczko LB. 2009. Molecular phylogeny of the Drosophila tripunctata and closely related species groups (Diptera: Drosophilidae). Mol. Phylogenet. Evol. 51:595–600.

Heed WB, Wheeler MR. 1957. Thirteen new species in the genus Drosophila from the Neotropical region. Univ. Texas Publ. 5721:17–38.

Huerta-Cepas J, Serra F, Bork P. 2016. ETE 3: Reconstruction, Analysis, and Visualization of Phylogenomic Data. Mol. Biol. Evol. 33:1635–1638.

Izumitani HF, Kusaka Y, Koshikawa S, Toda MJ, Katoh T. 2016. Phylogeography of the subgenus Drosophila (Diptera: Drosophilidae): Evolutionary history of faunal divergence between the old and the new worlds. PLoS One 11:e0160051.

Jeffroy O, Brinkmann H, Delsuc F, Philippe H. 2006. Phylogenomics: the beginning of incongruence? Trends Genet. 22:225–231.

Jeong S, Rebeiz M, Andolfatto P, Werner T, True J, Carroll SB. 2008. The Evolution of Gene Regulation Underlies a Morphological Difference between Two Drosophila Sister Species. Cell 132:783–793.

Kaneshiro KY. 1999. Sexual selection and speciation in Hawaiian Drosophila (Drosophilidae): A model system for research in Tephritidae. In: Fruit Flies (Tephritidae): Phylogeny and Evolution of Behavior.

Karageorgi M, Bräcker LB, Lebreton S, Minervino C, Cavey M, Siju KP, Grunwald Kadow IC, Gompel N, Prud’homme B. 2017. Evolution of Multiple Sensory Systems Drives Novel Egg-Laying Behavior in the Fruit Pest Drosophila suzukii. Curr. Biol. 27:847–853.

Katoh K, Standley DM. 2013. MAFFT multiple sequence alignment software version 7: Improvements in performance and usability. Mol. Biol. Evol. 30:772–780.

Katoh T, Izumitani HF, Yamashita S, Watada M. 2017. Multiple origins of Hawaiian drosophilids: Phylogeography of Scaptomyza Hardy (Diptera: Drosophilidae). Entomol. Sci. 20:33–44.

Katoh T, Nakaya D, Tamura K, Aotsuka T. 2007. Phylogeny of the Drosophila immigrans species group (Diptera: Drosophilidae) based on Adh and Gpdh sequences. Zoolog. Sci. 24:913–921.

Katoh T, Tamura K, Aotsuka T. 2000. Phylogenetic position of the subgenus Lordiphosa of the genus Drosophila (Diptera: Drosophilidae) inferred from alcohol dehydrogenase (Adh) gene sequences. J. Mol. Evol. 51:122–130.

Kellermann V, Van Heerwaarden B, Sgrò CM, Hoffmann AA. 2009. Fundamental evolutionary limits in ecological traits drive drosophila species distributions. Science 325:1244–1246.

Kim BY, Wang JR, Miller DE, Barmina O, Delaney E, Thompson A, Comeault AA, Peede D, D’Agostino ERR, Pelaez J, et al. 2020. Highly contiguous assemblies of 101 drosophilid genomes. bioRxiv.

Da Lage JL, Kergoat GJ, Maczkowiak F, Silvain JF, Cariou ML, Lachaise D. 2007. A phylogeny of Drosophilidae using the Amyrel gene: Questioning the Drosophila melanogaster species group boundaries. J. Zool. Syst. Evol. Res. 45:47–63.

Lang M, Murat S, Clark AG, Gouppil G, Blais C, Matzkin LM, Guittard É, Yoshiyama-Yanagawa T, Kataoka H, Niwa R, et al. 2012. Mutations in the neverland gene turned Drosophila pachea into an obligate specialist species. Science 337:1658–1661.

Lapoint RT, O’Grady PM, Whiteman NK. 2013. Diversification and dispersal of the Hawaiian Drosophilidae: The evolution of Scaptomyza. Mol. Phylogenet. Evol.

Lartillot N, Lepage T, Blanquart S. 2009. PhyloBayes 3: A Bayesian software package for phylogenetic reconstruction and molecular dating. Bioinformatics 25:2286–2288.

Lewis RL, Beckenbach AT, Mooers A. 2005. The phylogeny of the subgroups within the melanogaster species group: Likelihood tests on COI and COII sequences and a Bayesian estimate of phylogeny. Mol. Phylogenet. Evol. 37.

Van Der Linde K, Houle D, Spicer GS, Steppan SJ. 2010. A supermatrix-based molecular phylogeny of the family Drosophilidae. Genet. Res. (Camb). 92:25–38.

Máca J. 1992. Addition to the fauna of Drosophilidae, Camillidae, Curtonotidae, and Campichoetidae (Diptera) of Soviet Middle Asia. Annotationes Zoologicae et Botanicae 210: 1–8.

Longdon B, Hadfield JD, Day JP, Smith SCL, McGonigle JE, Cogni R, Cao C, Jiggins FM. 2015. The Causes and Consequences of Changes in Virulence following Pathogen Host Shifts. PLoS Pathog. 11:e1004728.

Magnacca KN, Foote D, O’Grady PM. 2008. A review of the endemic Hawaiian Drosophilidae and their host plants. Zootaxa 1728:1–58.

Malloch JR. 1934. Part VI. Diptera. In: Insects of Samoa. p. 267–312.

Markow T a., O’Grady P. 2006. Drosophila: A Guide to Species Identification and Use. Elsevier.

McGregor AP, Orgogozo V, Delon I, Zanet J, Srinivasan DG, Payre F, Stern DL. 2007. Morphological evolution through multiple cis-regulatory mutations at a single gene. Nature.

Mengual X, Kerr P, Norrbom AL, Barr NB, Lewis ML, Stapelfeldt AM, Scheffer SJ, Woods P, Islam MS, Korytkowski CA, et al. 2017. Phylogenetic relationships of the tribe Toxotrypanini (Diptera: Tephritidae) based on molecular characters. Mol. Phylogenet. Evol. 113:84–112.

Minh BQ, Hahn MW, Lanfear R. 2020. New methods to calculate concordance factors for phylogenomic datasets. Mol. Biol. Evol. 37:2727–2733.

Minh BQ, Schmidt HA, Chernomor O, Schrempf D, Woodhams MD, Von Haeseler A, Lanfear R, Teeling E. 2020. IQ-TREE 2: New Models and Efficient Methods for Phylogenetic Inference in the Genomic Era. Mol. Biol. Evol. 37:1530–1534.

Morales-Hojas R, Vieira J. 2012. Phylogenetic Patterns of Geographical and Ecological Diversification in the Subgenus Drosophila. PLoS One 7:e49552.

Morgan TH. 1910. Sex Limited Inheritance in Drosophila. Science 32:120–122.

O’Grady PM, Clark JB, Kidwell MG. 1998. Phylogeny of the Drosophila saltans species group based on combined analysis of nuclear and mitochondrial DNA sequences. Mol. Biol. Evol. 15:656–664.

O’Grady PM, DeSalle R. 2018. Phylogeny of the genus Drosophila. Genetics 209:1–25.

O’Grady PM, Lapoint RT, Bonacum J, Lasola J, Owen E, Wu Y, DeSalle R. 2011. Phylogenetic and ecological relationships of the Hawaiian Drosophila inferred by mitochondrial DNA analysis. Mol. Phylogenet. Evol.

O’Grady PM, Magnacca K, Lapoint RT. 2009. Drosophila. In: Gillespie R, Clague D, editors. Encyclopedia of Islands. University of California press, Berkeley, CA. p. 232–235.

Okada T. 1967. A revision of the subgenus Hirtodrosophila of the Old World, with descriptions of some new species and subspecies (Diptera, Drosophilidae, Drosophila). Mushi 41:1–36.

Okada T. 1984. The Genus Collessia of Japan (Diptera: Drosophilidae). Proc. Japanese Soc. Syst. Zool. 29:57–58.

Okada T. 1988. Family Drosophilidae (Diptera) from the Lund University Ceylon Expedition in 1962 and Borneo collections in 1978-1979. Entomol. Scand. 30:109–149.

Peluffo AE, Nuez I, Debat V, Savisaar R, Stern DL, Orgogozo V. 2015. A major locus controls a genital shape difference involved in reproductive isolation between Drosophila yakuba and Drosophila santomea. G3 Genes, Genomes, Genet. 5:2893–2901.

Philippe H, Sörhannus U, Baroin A, Perasso R, Gasse F, Adoutte A. 1994. Comparison of molecular and paleontological data in diatoms suggests a major gap in the fossil record. J. Evol. Biol. 7:247–265.

Prigent SR, Le Gall P, Mbunda SW, Veuille M. 2013. Seasonal and altitudinal structure of drosophilid communities on Mt Oku (Cameroon volcanic line). Comptes Rendus - Geosci. 345:316–326.

Prigent SR, Suwalski A, Veuille M. 2017. Connecting systematic and ecological studies using DNA barcoding in a population survey of Drosophilidae (Diptera) from Mt Oku (Cameroon). Eur. J. Taxon. 2017.

Remsen J, O’Grady P. 2002. Phylogeny of Drosophilinae (Diptera: Drosophilidae), with comments on combined analysis and character support. Mol. Phylogenet. Evol. 24.

Robe L. J., Cordeiro J, Loreto ELS, Valente VLS. 2010. Taxonomic boundaries, phylogenetic relationships and biogeography of the Drosophila willistoni subgroup (Diptera: Drosophilidae). Genetica 138.

Robe Lizandra J., Loreto ELS, Valente VLS. 2010. Radiation of the, Drosophila” subgenus (Drosophilidae, Diptera) in the Neotropics. J. Zool. Syst. Evol. Res. 48:310–321.

Robe LJ, Valente VLS, Budnik M, Loreto ÉLS. 2005. Molecular phylogeny of the subgenus Drosophila (Diptera, Drosophilidae) with an emphasis on Neotropical species and groups: A nuclear versus mitochondrial gene approach. Mol. Phylogenet. Evol. 36:623–640.

Robe Lizandra J., Valente VLS, Loreto ELS. 2010. Phylogenetic relationships and macro-evolutionary patterns within the Drosophila tripunctata “radiation” (Diptera: Drosophilidae). Genetica 138:725–735.

Russo CAM, Mello B, Frazão A, Voloch CM. 2013. Phylogenetic analysis and a time tree for a large drosophilid data set (Diptera: Drosophilidae). Zool. J. Linn. Soc. 169:765–775.

Sackton TB, Lazzaro BP, Schlenke TA, Evans JD, Hultmark D, Clark AG. 2007. Dynamic evolution of the innate immune system in Drosophila. Nat. Genet. 39:1461–1468.

Sigurdsen T, Green DM. 2011. The origin of modern amphibians: A re-evaluation. Zool. J. Linn. Soc. 162:457–469.

Stamatakis A. 2014. RAxML version 8: A tool for phylogenetic analysis and post-nalysis of large phylogenies. Bioinformatics 30:1312–1313.

Stark JB, O’Grady PM. 2010. Morphological variation in the forelegs of the Hawaiian Drosophilidae. I. The AMC clade. J. Morphol. 271:86–103.

Sturtevant A. H. 1959. Thomas Hunt Morgan. In: A biographical memoir of national academy of sciences. Vol. 33. p. 283–325.

Sucena E, Delon I, Jones I, Payre F, Stern DL. 2003. Regulatory evolution of shavenbaby/ovo underlies multiple cases of morphological parallelism. Nature 424:935–938.

Tanaka K, Barmina O, Kopp A. 2009. Distinct developmental mechanisms underlie the evolutionary diversification of Drosophila sex combs. Proc. Natl. Acad. Sci. U. S. A. 106:4764–4769.

Tatarenkov A, Zurovcová M, Ayala FJ. 2001. Ddc and amd sequences resolve phylogenetic relationships of Drosophila [3]. Mol. Phylogenet. Evol. 20:321–325.

TaxoDros. 2020. TaxoDros, the database on Taxonomy of Drosophilidae.

Throckmorton L. 1962. The problem of phylogeny in the genus Drosophila. Univ. Texas Publ. 2:207–343.

Throckmorton L. 1975. The phylogeny, ecology and geography of Drosophila. In: King R, editor. Handbook of genetics. New York. p. 421–469.

Throckmorton L. 1982. Pathways of evolution in the genus Drosophila and the founding of the repleta group. In: Barker J, Starmer W, editors. Ecological Genetics and Evolution: the Cactus-Yeast-Drosophila Model System. Academic Press, New York. p. 33–47.

Trinder M, Daisley BA, Dube JS, Reid G. 2017. Drosophila melanogaster as a high-throughput model for host-microbiota interactions. Front. Microbiol. 8:751.

Tsacas L. 2002. Le nouveau complexe africain Drosophila loiciana et l’espèce apparentée D. matileana n. sp. (Diptera: Drosophilidae). Ann. la Société Entomol. Fr. 38:57–70.

Tsacas L, Chassagnard M-T. 1992. Les relations Araceae-Drosophilidae. Drosophila aracea une espèce anthophile associée à l’aracée Xanthosoma robustum au Mexique (Diptera: Drosophilidae). Ann. la Société Entomol. Fr. 28:421–439.

Ugur B, Chen K, Bellen HJ. 2016. Drosophila tools and assays for the study of human diseases. Dis. Model. Mech. 9:235–244.

Wakahama K-I, Shinohara T, Hatsumi M, Uchida S, Kitagawa O. 1983. Metaphase chromosome configuration of the immgrans species group of Drosophila. Japanese J. Genet. 57:315–326.

Wartlick O, Mumcu P, Jülicher F, Gonzalez-Gaitan M. 2011. Understanding morphogenetic growth control - lessons from flies. Nat. Rev. Mol. Cell Biol. 12:594–604.

Wheeler MR. 1959. A Nomenclatural Study of the Genus Drosophila. Univ. Texas Publ. 5914:181–205.

Wheeler MR, Kambysellis MP. 1966. Notes on the Drosophilidae (Diptera) of Samoa. Univ. Texas Publ. 6615.

Wiegmann BM, Richards S. 2018. Genomes of Diptera. Curr. Opin. Insect Sci. 25:116–124.

Wiegmann BM, Trautwein MD, Winkler IS, Barr NB, Kim J-W, Lambkin C, Bertone M a, Cassel BK, Bayless KM, Heimberg AM, et al. 2011. Episodic radiations in the fly tree of life. Proc. Natl. Acad. Sci. U. S. A. 108:5690–5695.

Yassin A. 2013. Phylogenetic classification of the Drosophilidae Rondani (Diptera): The role of morphology in the postgenomic era. Syst. Entomol. 38:349–364.

Yassin A, Debat V, Bastide H, Gidaszewski N, David JR, Pool JE. 2016. Recurrent specialization on a toxic fruit in an island Drosophila population. Proc. Natl. Acad. Sci. U. S. A. 113:4771–4776.

Yassin A, Delaney EK, Reddiex AJ, Seher TD, Bastide H, Appleton NC, Lack JB, David JR, Chenoweth SF, Pool JE, et al. 2016. The pdm3 Locus Is a Hotspot for Recurrent Evolution of Female-Limited Color Dimorphism in Drosophila. Curr. Biol. 26:2412–2422.

Yassin A, Da Lage J-L, David JR, Kondo M, Madi-Ravazzi L, Prigent SR, Toda MJ. 2010. Polyphyly of the Zaprionus genus group (Diptera: Drosophilidae). Mol. Phylogenet. Evol. 55:335–339.

Zhang W, Toda MJ. 1992. A new species-subgroup of the Drosophila immigrans species group (Diptera, Drosophilidae) with description of two new species from China and revision of taxonomic terminology. Japanese J. Entomol. 60:839–850.

Zhang Y, Izumitani HF, Katoh TK, Finet C, Toda MJ, Watabe H, Katoh Toru. 2021. Phylogeny and evolution of mycophagy in the Zygothrica genus group (Diptera: Drosophilidae).

