## Supplementary figures and images for "DrosoPhyla: genomic resources for drosophilid phylogeny and systematics"

### Figure S1

#### IQ-TREE

##### GTR+R+FO

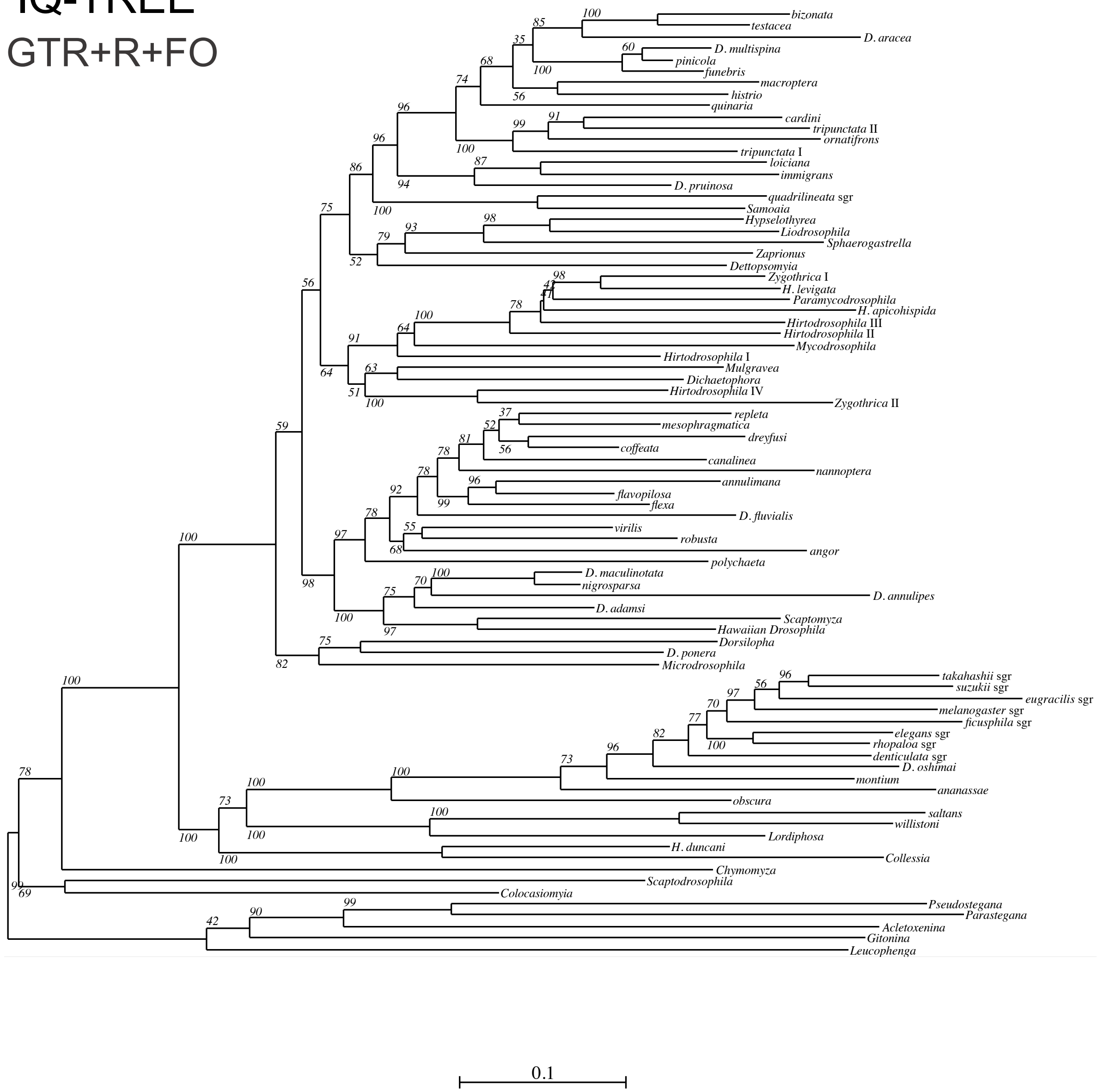


#### PhyloBayes

##### GTR+Γ

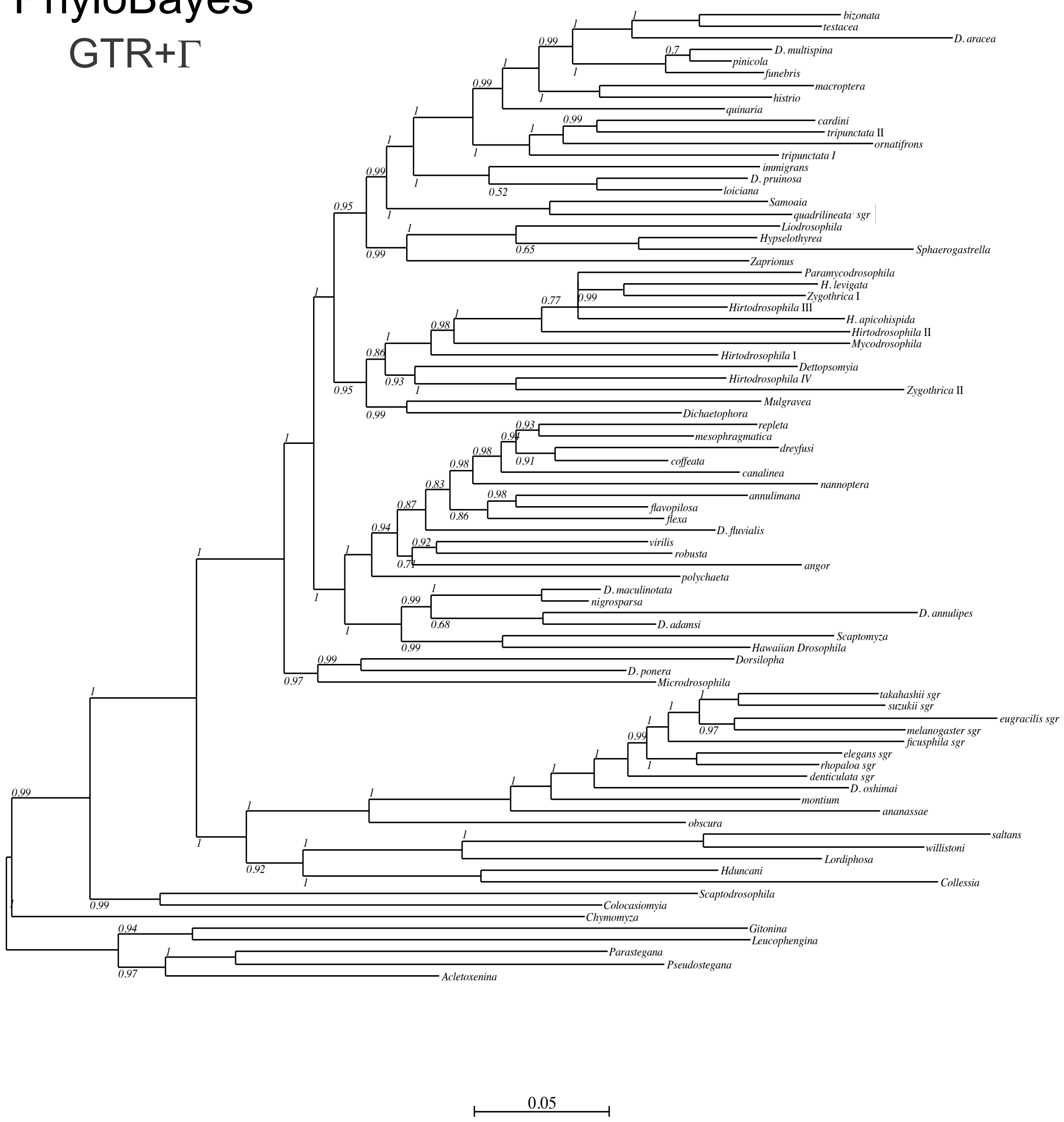

### Figure S2

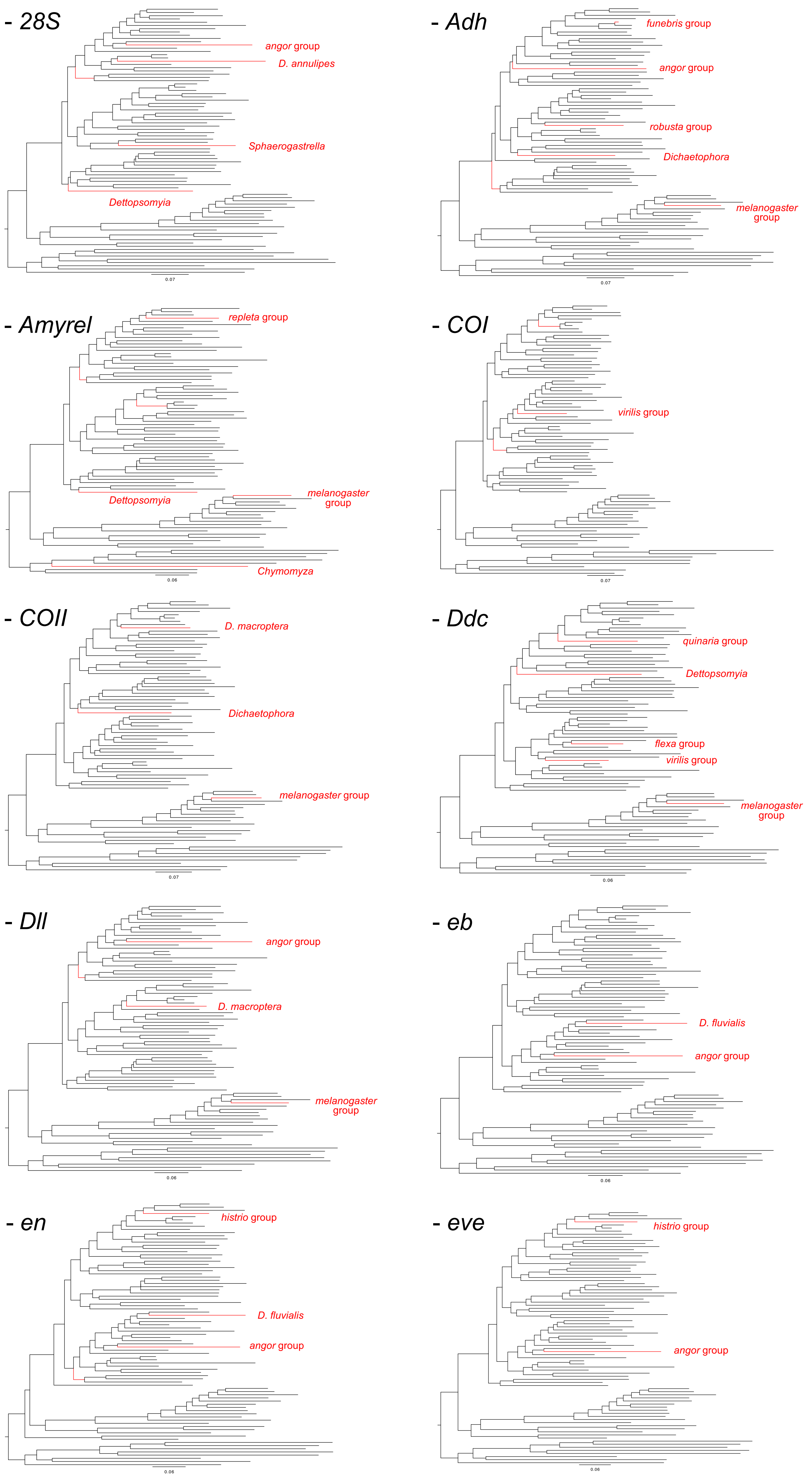

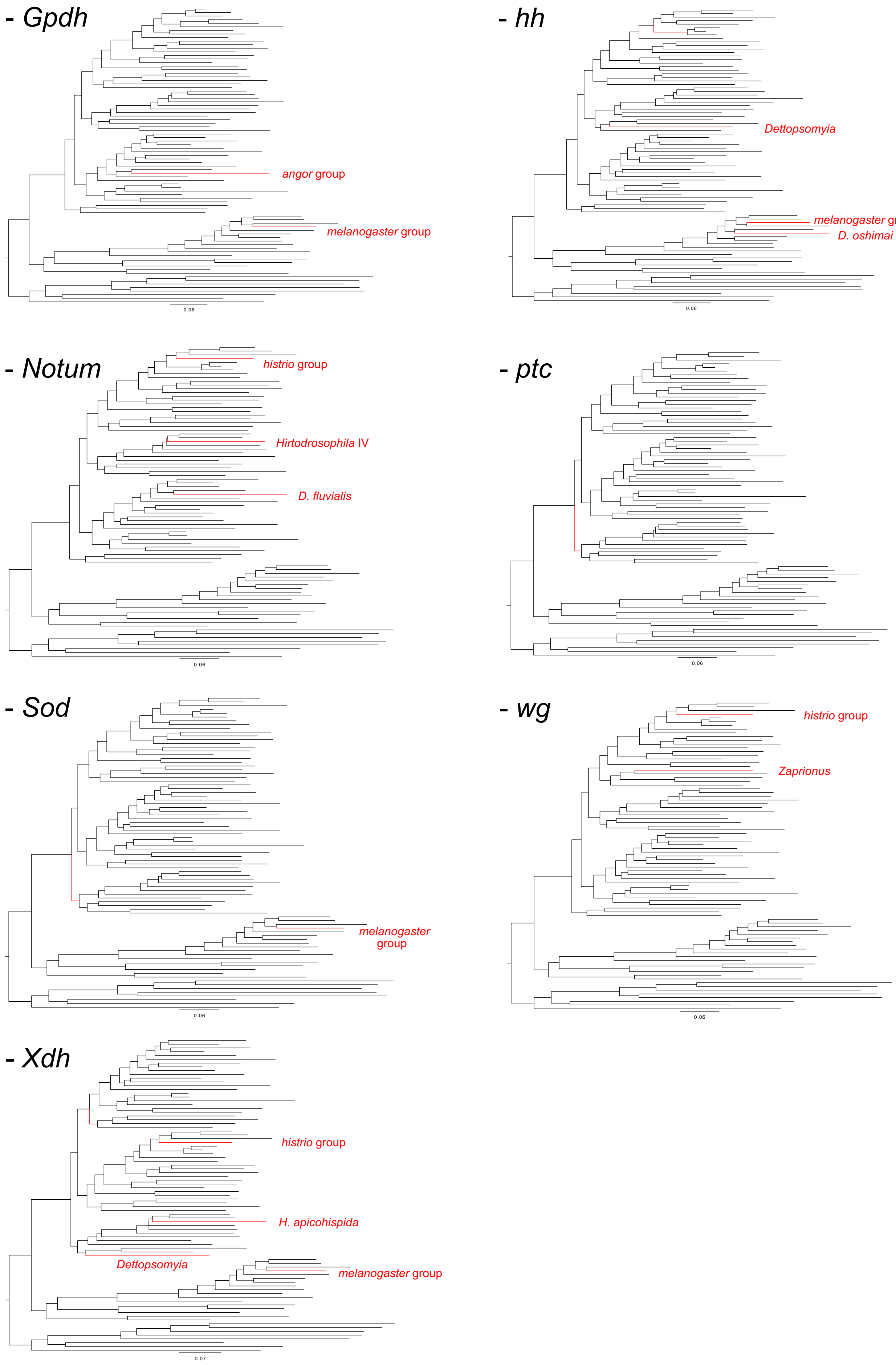

### Figure S3

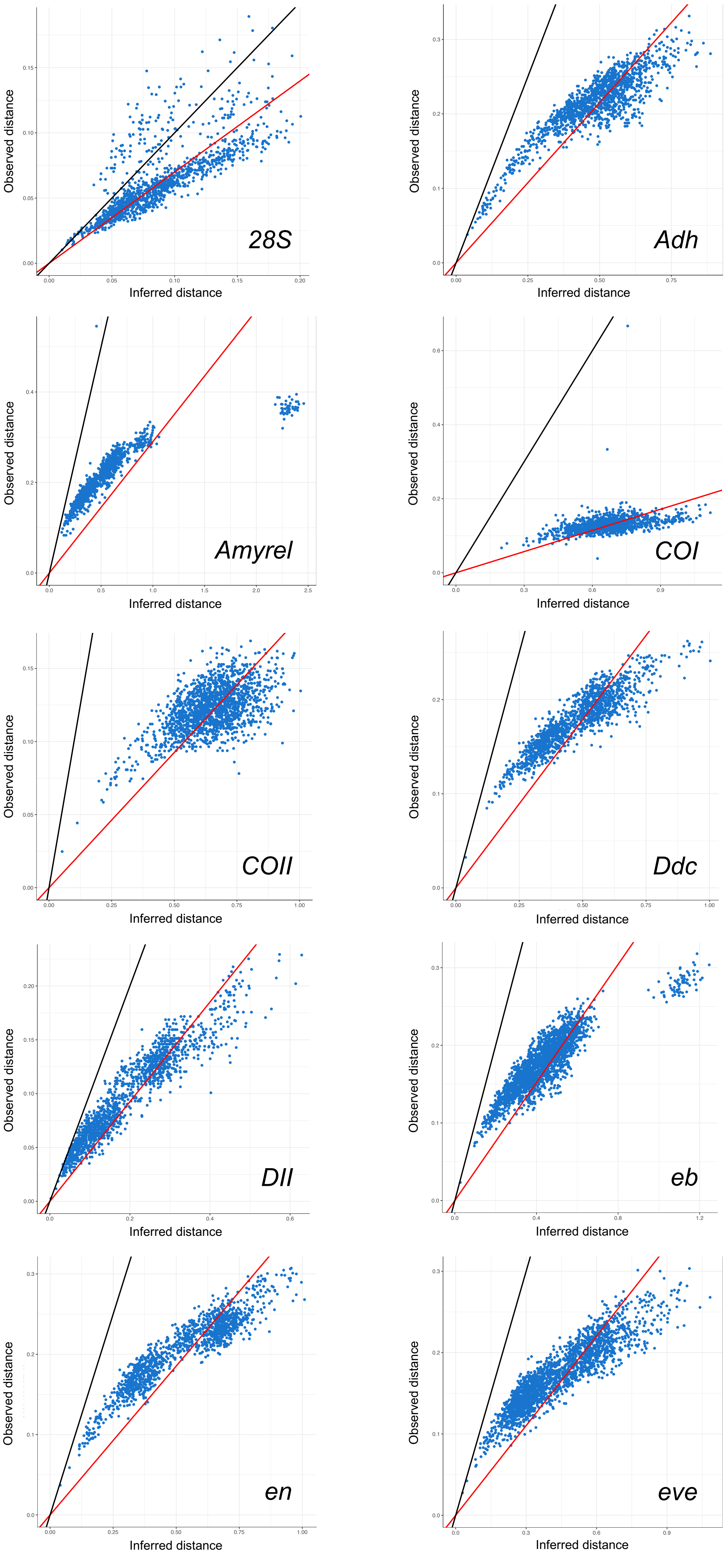

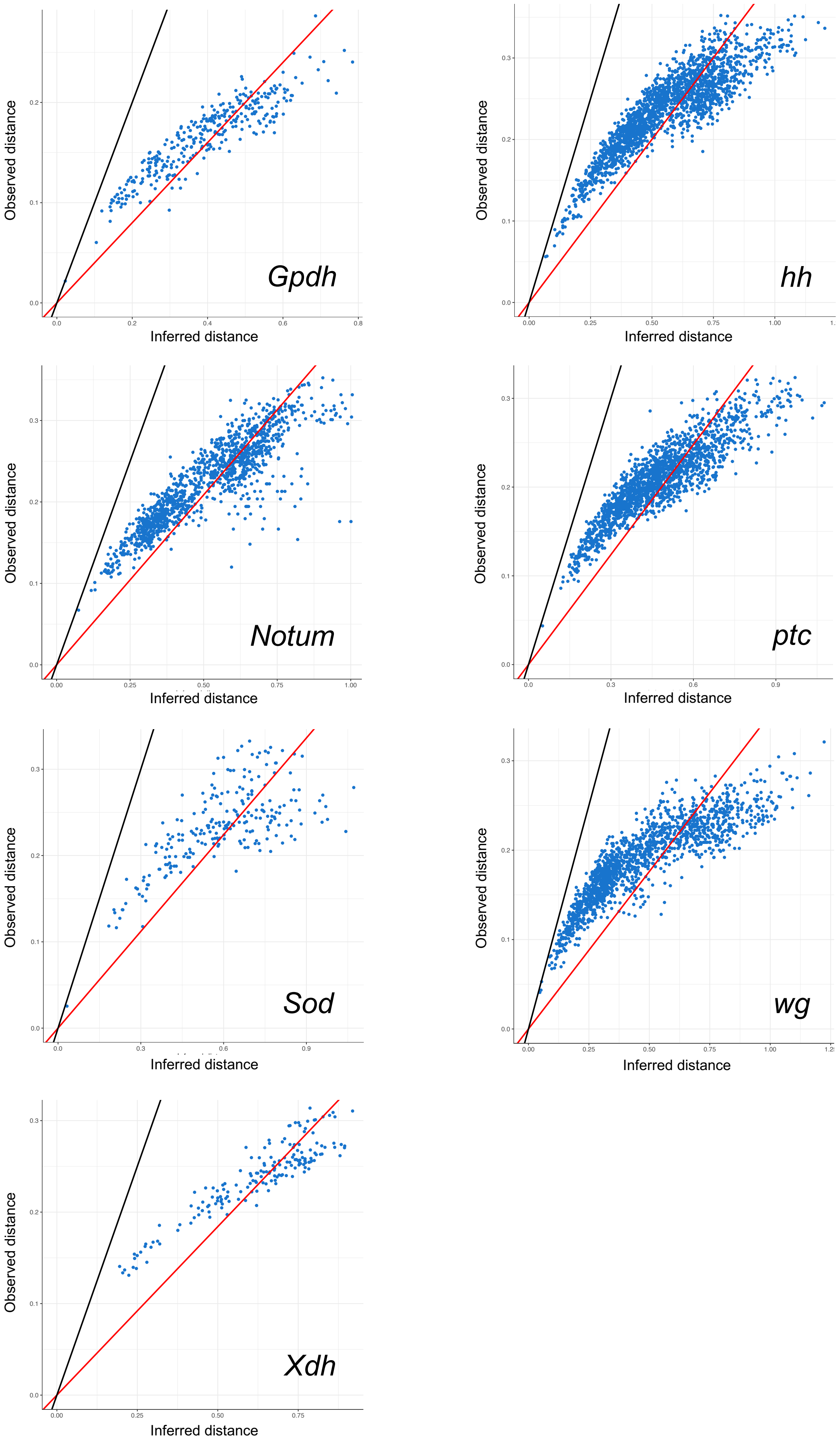
